# What can astrocytes compute?

**DOI:** 10.1101/2021.10.20.465192

**Authors:** Erik J. Peterson

**Affiliations:** Department of Psychology, Carnegie Mellon University

## Abstract

A foundational result in neural computation was proving the firing-rate model of neurons defines a universal function approximator. In this paper I prove it is possible for astrocytes to compute anything neurons can, by extending this original proof to a model of calcium waves in astrocytes. I confirm this in a series of computer simulations. The major limit for astrocytes, it turns out, is not their ability to learn solutions but the computational complexity of their network. I suggest some initial experiments that might be used to confirm these predictions.

---

We’ve known for sometime astrocytes generate elaborate calcium waves, and that somehow these waves are important for learning [40, 16]. But two recent experiments offer an intriguing new possibility for computation in these cells. Mu et al (2019) showed direct signal integration by astrocytes leading to motor control in zebrafish [34]. Slezack et al (2019) showed direct integration of visual information, and behavioral state, in mice [41]. While it is was well known that astrocytes generate calcium waves [29, 19, 11], their role was thought to be limited to tuning neurons as part of the tripartite synapse [5, 45, 36]. These new results from [34] and [41], along with older speculations [37], however suggest a direct role for astrocytes in computation.

Inspired by these experimental results the aim of this paper is to use theoretical analysis to “rule in” a large range of computations as mathematically possible for astrocytes, reaching well beyond what we have evidence for experimentally.

I consider three questions.

1. Can the fundamental universal approximator proofs for neurons be extended to astrocytes?
2. If astrocytes can approximate universally, why do neurons exist?
3. Can simulations of astrocytes learn as well as a simulations of neurons?

## Results

The results have three parts answering the three questions from the introduction. First is a theoretical study of function approximation in astrocytes. Second, I analyze the computational complexity of astrocyte networks. Third is some computer simulations.

### Neural approximation

Let’s first review biological neural networks reduced to firing-rate models of point cells. From there we can define a universal approximation.

The kind of two-layer neural network we are interested in is shown in Figure 1a and has the mathematical form,

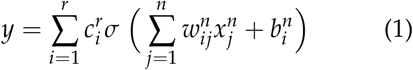

**Figure 1:**
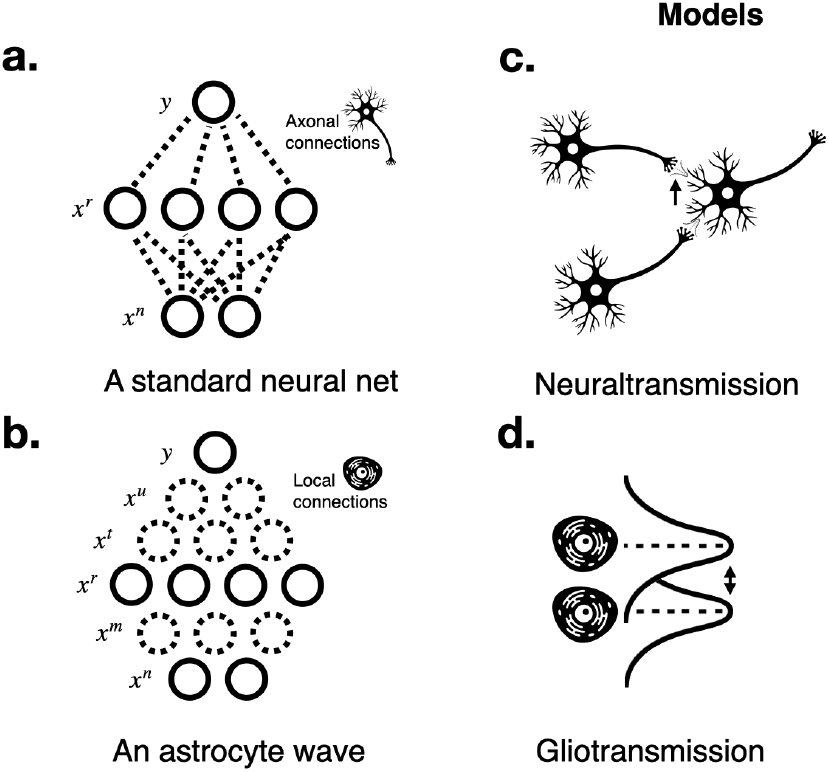
Models. **a**-**b** Diagram of communication in firing rate neural network (wij*a*), and an astrocyte wave with *k* = 2 neighbors (*b*). **c**-**d** Modes of transmission. Neuronal transmission (*a*) generally occurs in a well defined synapse, which among other things tightly controls the diffusion of the transmitter. Gliotransmission is far less confined, and can diffuse some distance leading to a potential for cross-talk between cells, a.k.a. leak.

Where the input 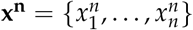 is a vector of size *n*, each *x_i_* ∈ **R**, and the output *y* is a single real value *y* ∈ **R**. Here *w_ij_* ∈ **R** is the weight between ith element of the input and the jth unit in layer *n*, 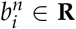 is the threshold of the ith unit, and 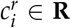 is the weight between the ith unit in layer *r* (which is also the output *y*). I use *σ* to denote some nonlinear function, restricted as described below.

Note the use of subscripts to denote indexing under summation and superscripts to denote the layer each term belongs to. In a two layer neural network these superscripts seem redundant. But as we move to study many-layered astrocyte wave networks they are a convenient way to prevent proliferation of excess terms.

It is known that two-layer neural networks [2] are universal function approximators. All approximator proofs follow the same form. They remake the question of how good an approximation is into a question about the “density” between two sets of numbers [38]. Density formalized as requiring every element in *M* is within a neighborhood *ε* > 0 of an element in *C*. If this criterion holds, then *M* is said to be dense in *C*. Thiss kind of density is an abstract way to measure near equivalence in sets, and as a result approximation between functions.

Our target function I denote *f*, and set up as a continuous real function *f ∈ C*(ℝ^*N*^). It is this function we wish to approximate. To do that we’ll be using an approximator *F*, given in this case by Equation 1.

To ask density questions about these we’ll need sets, not function. To that end, it is common to focus on some compact subset *K* of ℝ^*n*^. The set for *f* is then {*f* (*x*) : *x ∈ K*}. For the approximator *F* things are slightly complex, because we need a parameterized set. To get that I follow [38] and define a set building scheme, *M_r_* (below), otherwise known as the spanning set.

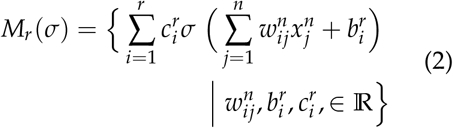

Armed with a way to build sets for *F* and *f*, on the “ball” *K*, we can then ask the following question. For which *σ* is it true that,

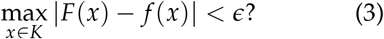

Answering this question positively means proving universal approximation. For neural networks of the kind above this has been studied for several kinds of *σ*, but also for may other networks beyond two-layers [12, 26, 2, 38, 44, 31, 33, 44, 23, 32, 28]. In this work, I borrow a classic theorem for the two-layer network as given by [38]. It is modified to include a continuous nonzero derivative on *σ*.

#### Theorem 0.1 (Neural approximation)

*Let σ by any nonlinear function that is not a polynomial function and has a continuous nonzero derivative at some point. Then finite sums of the form in Equation 1 are dense in C* (ℝ^*n*^) *on K. In other words, given any f* (*x*) ∈ *C*(ℝ^*n*^) *and ε >* 0 *there is function F* (*x*) *for which,* |*F*(*x*) − *f* (*x*)| > *ε*, *for all x* ∈ *K.*

Proving approximation only means *F* can “do the job” approximating *f*. It does not ensure an efficient method to cause *F* to approximate *f* exists or is known. Another way to explain this limit is to say that density proofs are not constructive proofs. I take up construction later on.

### Astrocyte equivalence

The model of astrocytes I’ll consider is shown in Figure 1b. Compare it to a standard two-layer neural network, in Figure 1a. I assume,

- Astrocyte communication is only between *k* nearest neighbors in the “forward” direction. (Forward is defined as the flow from input to output).
- Astrocytes can be modeled as point cells on an ordered rectangular grid.
- That gliotransmission is the dominant mode of communication.
- (If instead gap junctions are dominant this would improve the specificity of communications and so improve practical approximation performance. See the Astrocyte complexity section below.)
- The lack of synapses means transmitters are released from the cell wall uniformly, and “leaks” across to neighborhoods. (We show this leak in Figure 1d as a Gaussian; Contrast this leak to the “exact” synapses in a neural network as shown in Figure 1c.)
- This leak does not extend past *k* neighbors.

The limited connectivity between astrocytes in the model means writing down the general equation for astrocytes is more complex than for neurons. I show an example of a *r* = 4, *k* = 2 astrocytes network in Figure 1b and mathematically below. This example matches the max width *r* = 4 of the two layer neural network depicted in Figure 1a.

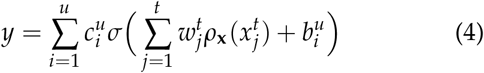

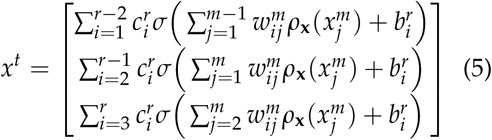

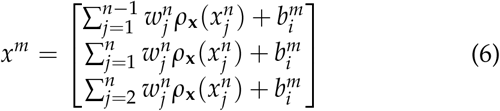

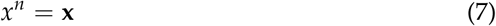

I assume an astrocyte network should take the general form shown in Equation 4, generalized for any choice of finite *n* and *r > n* + 1 [28], where *r* denotes the maximum width of the widest layer. The function *ρ*_x_ stands in for a generic convolution operation take on each element in some *x* based in part on the vector x, where refers to some layer input. I other words, this is how I model transmitter diffusion, or “leak”. Having introduced the physical idea of leak for now though I will neglect it, by setting *ρ*_x_ to 1, and treating astrocytes as if they formed direct connections. Justifications for, and tests of, these assumptions are found in the simulations below. With these we can now prove any astrocyte network *G* can be made equivalent to any two-layer neural network *F*, which is itself a universal approximator. Without loss of generality I write the spanning set for astrocytes *P*(*σ*), as a series of sum and compositions based on Equations 4, generalizable as necessary to other *n* and *r*.

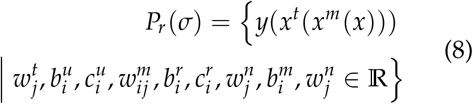

#### Theorem 0.2 (Astrocyte equivalence)

*Let σ by any nonlinear function that is not a polynomial function and has a continuous nonzero derivative at some point. Then generalized finite sums of the form in Equation 4 can be made equivalent with the form in Equation 1 there if F* (*x*) *is dense then G* (*x*) *is also dense in C*(ℝ^*N*^) *on K. In other words, given any g* ∈ *P_r_*(*σ*) *there is a neural network F* ∈ *M*(*σ*) *for which* |*G*(*x*) − *F*(*x*)| = 0, *for allx* ∈ *K. And therefore given any f* (*x*) ∈ *C*(ℝ^*N*^) *and ε >* 0 *there is function G*(*x*) *for which,* |*G*(*x*) − *f* (*x*)| < *ε for all x ∈ K*

The proof for this is trivial and proceeds in two parts. We first recognize the inner sum in Equation 1, that is 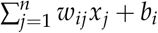, is an simply an affine function, which can perfectly reconstructed by linearizing the corresponding region in *G*. That is, layers *n* through *r* are linearized allowing for an exact superposition solution.

Each layer and set of connection in the “outer” sum, 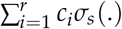 is composed of small networks mathematically equivalent to neural networks with *n* = 3, and is therefore composed of compositions of nonlinear terms that are already proven to be universal approximators. In other words, the outer sum is necessarily step-wise dense in *C* by reusing the argument made for neurons.

This proof is not intended to be a guide to constructing astrocyte networks. Nor is the linearizing step necessary for construction of working waves, as I show in the next section. This theoretical approach is instead a means to an end to establish density.

### Astrocyte complexity

Before giving mathematical details the big picture for the complexity of astrocyte waves is easy to state. To span distances astrocytes must pass messages between cells. Each cell is in a sense a new set of parameters, and so complexity of the whole model increases, compared to neurons.

If we assume neurons axons can span any distance |*m − n*|, then for a neuronal network we can move from *m* to *n* in *l* = 1 layers, and a total of *m* + *n* cells.

The lack of axons on astrocytes means however there is a necessary link between the size or width of a feed forward network and it’s depth. If each cell rests on the corners of a regular grid, going from from input size *m* to output size *n*, where *m* ≠ *n*, it will take 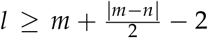 layer steps, if both *m* and *n* are even or odd. It will take 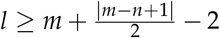 steps if there is *m* is even and *n* odd, or vice versa. If the widths are the same, so *m* = *n*, the layer “penalty” is *l* = *m* − 2. If *l* is the difference in layer numbers between astrocytes and neurons is *l*, then the cell number penalty is the sum over *l*, with the generic term *o_i_* standing for the width of each layer. That is, 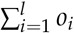.

In computer science terms an O(*l*), or linear penalty, for a computation is still viable, even efficient, computation. I argue this viability is not necessarily so in biology. For example, imagine neurons had not been “invented” and circuits were limited to wave/step computations and we continue to use grid models to make the calculations convenient. If the wave transmission in a single astrocyte takes *t_a_* seconds, the the total transmission time is *T_a_* = *t_a_* * *l* for astrocytes. It would be *t_a_* for neurons. If we modestly assume a very small neural circuit to compare to, say *m* = 2 and *n* = 12 so *l* = 4, then *T_a_* has grown four fold. Consider what this means for motor output, as an example. If the best motor reaction time of an animal with neurons is 10 ms, it would grow to 40 ms with astrocytes. This is a substantial difference in reaction time for say a prey escaping an agile predator. Now consider what a linear time penalty would imply for mammalian brains with their trillions of connections [1, 35].

In sum, the analysis in this paper implies the neuron’s innovation—with the introduction of axons and synapses—-was not the ability to learn. It was the ability to compute as efficiently, in terms of width, depth, and cell number. But this is not the whole story, perhaps. Our view has been looking at what it would take to make astrocytes mimic neurons, exactly. In some ways this provides insight into the base question of “why neurons”?. But, there is no reason for evolution to hold this perspective, exactly.

Astrocytes can be expected to perform more efficiently than neurons when there is some “sympathy” between them and the learning problem at hand [50]. For example, in [34] et al astrocytes act to integrate incoming signals. If this integration was done instead using neurons, it would require several cells, or circuits, with exacting properties [48, 8, 15]. Meanwhile slow calcium dynamics, combined with connectionist waves, generate natural integration [42, 14, 34, 41]. That is despite the analysis above, we have meaningful evidence that astrocytes can be more efficient in terms of cell number, for some computations which match their properties like integration. The question I pose is, given astrocytes can act as universal approximators, what other functions might they fulfil efficiently which have been so far missed in our experiments?

### Simulations

Astrocytes learn (nearly) as well as neurons in three test simulations. Both astrocyte and neural models were trained by stochastic gradient descent, using identical procedures and parameters. The learning tasks were one classic nonlinear learning test (the XOR problem) and two classic visual recognition datasets (MINST digits and fashion). These tasks are shown in Figure 2.

**Figure 2:**
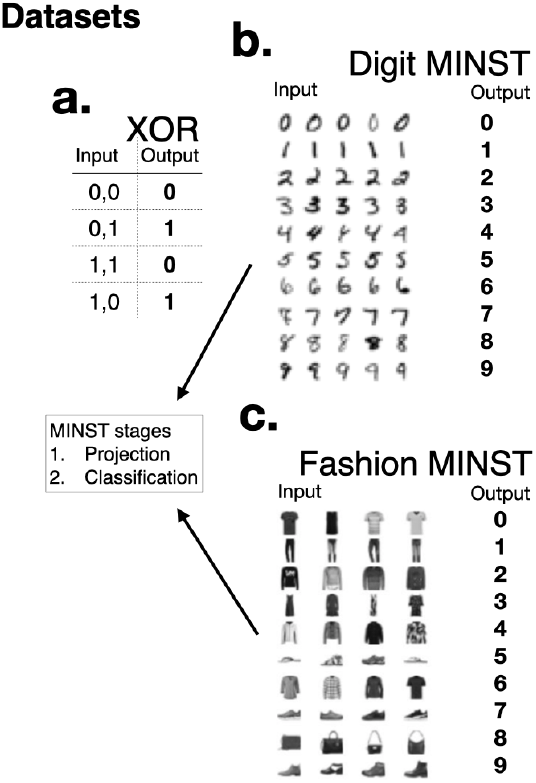
**a**-**b**. Datasets. We considered three decision tasks. **a.** XOR task. **b.** MINST digit recognition **c.** MINST fashion recognition. Learning in MINST sets had two distinct stages (grey box). First the raw pixel input data had its dimensionality reduced to 1d vector (*z* = 20). This representation was used as the input to the learned classifier networks, both neurons and astrocytes.

To design astrocytes waves I intermingled the three basic courses a feed-forward wave can take–spreading themselves out, collapsing or gathering themselves in, or simply sliding forward with no change in width. The final model architectures are shown in *Table 1*, and *2* (Methods).

**Table 1:** Astrocyte model and training hyper-parameters. Note: *s* denotes spread steps, *g* denotes gather, and *d* denotes slide. The learning rate for the XOR network was 0.001. The two MINST models have a learning rate of 0.004, and a 128 image batch size.

| Network | Layers | Input size | Output size | Layer steps |
| --- | --- | --- | --- | --- |
| XOR | 13 | 2 | 1 | (s,d,s,d,s,d,g,s,g,s,g,g,g) |
| Digits | 10 | 20 | 10 | (g,s,g,s,g,s,g,s,g,s,g) |
| Fashion | 10 | 20 | 10 | (g,s,g,s,g,s,g,s,g,s,g) |

**Table 2:** Network models and training hyper-parameters. The learning rate for the XOR network was 0.001. The two MINST models have a learning rate of 0.004, and a 128 image batch size.

| Network | Layers | Input size | Output size | Hidden sizes |
| --- | --- | --- | --- | --- |
| XOR | 1 | 2 | 1 | 1 |
| Digits | 1 | 20 | 10 | 20 |
| Fashion | 1 | 20 | 10 | 20 |

The change in size from input image to output class was too large, given astrocytes complexity. And the various tricks for fixing vanishing gradients available to the machine learning practitioner do not seem biologically sound. To overcome the complexity limits of astrocytes, and the vanishing gradients which follow from it, all vision tasks had two stages. First the high dimensional images were projected to a low dimensional space. This was done with neural negotiation. One approach mimicked the fixed random sparse connections sometimes reported in cortex [6, 39]. The other approach trained a variational autoencoder [30] during astrocyte training, in an online way. This was termed co-learning, and it mimicked a learning process where astrocytes adjust online to ongoing changes/learning in their (presumptively) neuronal input.

On the XOR task astrocytes and neurons showed perfect accuracy (Figure 3**a**). In both MINST tasks astrocyte performance was reduced between 0.05 and 0.08 (3**b-e**). Variance was also substantially higher in astrocytes and they were substantially slower to train (see inset panels in 3). The difference in speed of learning was as predicted. See *Complexity*

**Figure 3:**
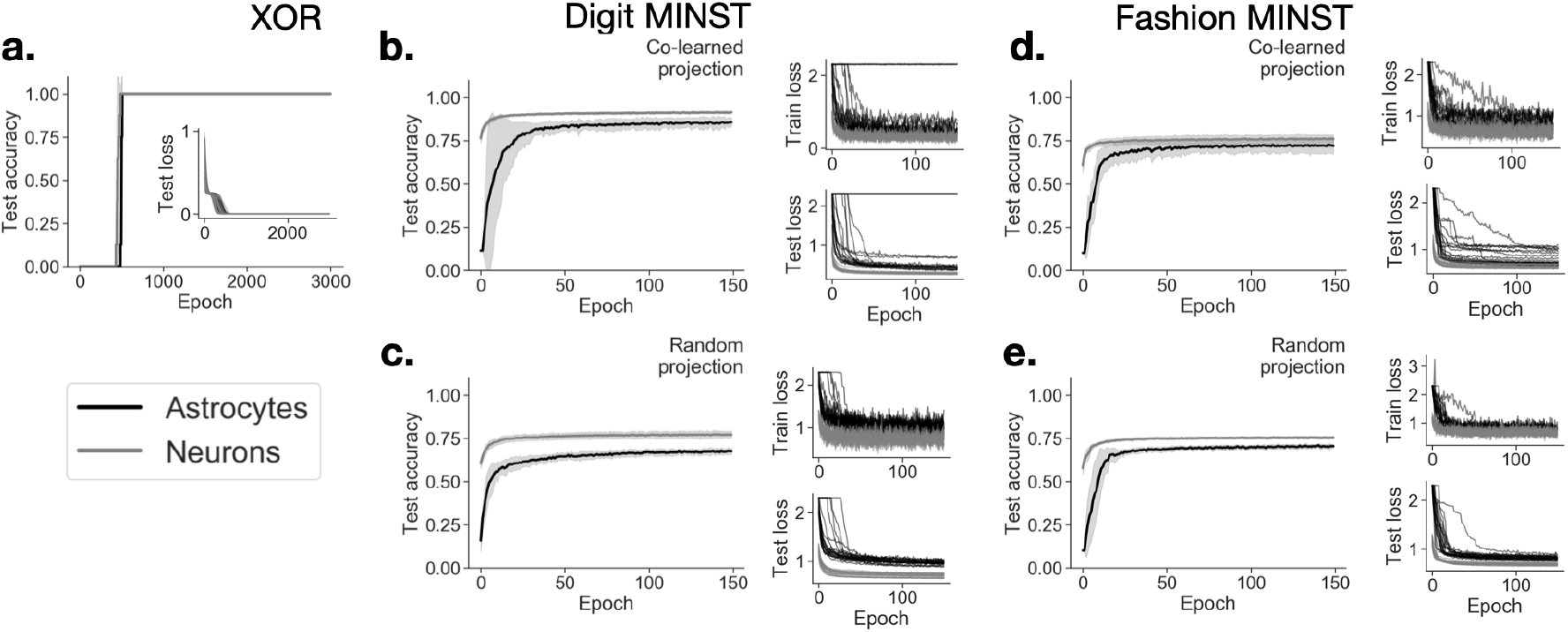
Learning performance. **a**. XOR task classification accuracy as a function of training epoch, on the test set (which is the training set for this task). Inset plot is individual loss curves for all 20 experiments. **b**-**c**. MINST digit classification accuracy for co-learning and random projection schemes. To the right of each are the individual train/text loss curves. **d**-**e**. MINST fashion classification accuracy for co-learning and random projection schemes. To the right of each are the individual train/text loss curves.

Connections in this working model were limited to each cell’s *k* = 3 nearest downstream neighbors. However in this model I was free to explore transmitter leak between greater than *k* neighbors, modeled by a 1d Gaussian convolution of each layers output. These simulations also let me explore biological factors like additive noise, and communication failures (dropout) between neighbors (*Methods*). See, Figure 4.

**Figure 4:**
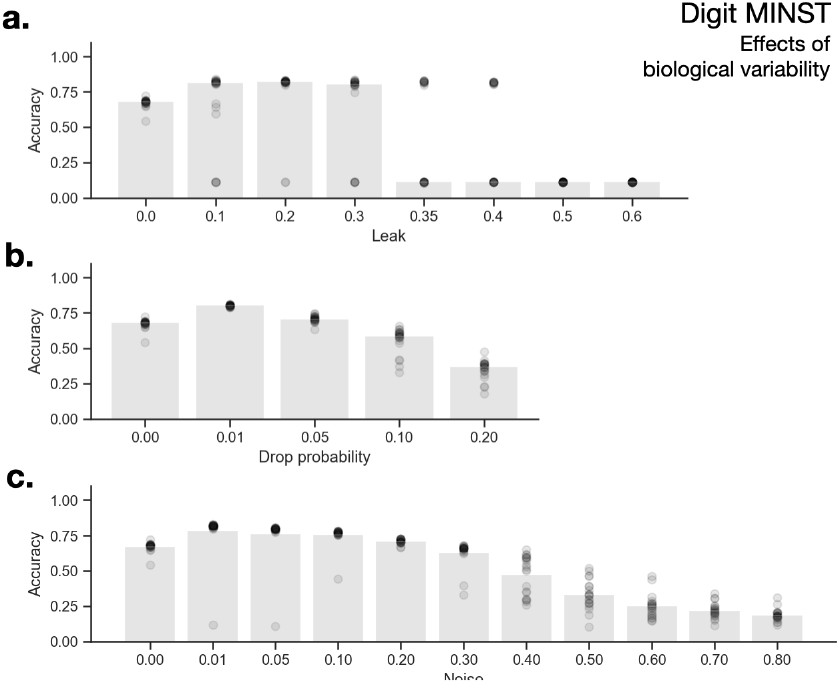
Biological realism. Leak (diffusion), communication failures (message dropping), and noise (message corruption) and their effects on astrocyte learning performance. **a.** Final test accuracy after convolution with a spatial Gaussian kernel (0-0.6 standard deviations). **b.** Final test accuracy with probabilistic connection loss between neighbors (0.0-0.20). **c.** Final test accuracy with connection noise (0.0-0.8 standard deviations). Points represent 20 individual experiments. Grey bars represent median accuracy.

Performance is robust to <0.3 standard deviations of leak, followed by sharp decline. In contrast, injected (additive) transmission noise, and signal loss, had much smoother declines in classification. The biological significance of these patterns is unclear. Overall these biological perturbations showed what seems to be reasonable robustness leak, noise, and signal loss. There is however a significant caveat. It is difficult to know how plausible the magnitude of these perturbations were compared to what real astrocytes, biological systems, endure. Detailed data does at present seem to be available.

## Discussion

### Limtations

I have studied a highly simplified model of feed-forward computations, with nearest neighbor connections between cells. I presume this is a minimal but adequate account of calcium waves. That said, this simplified model neglects:

- Lateral and recurrent calcium dynamics [20, 40, 49].
- Gap junctions and their associated plasticity [16].
- Neuronal-astrocyte interactions during wave-computation [21, 40].
- Calcium microdomains in individual cells. The complex, sometimes gated, calcium-gliotransmitter relationship [4].
- Astrocytes are a diverse cell type, with diverse shapes roles and electrical properties [27]

I think of these details as adding more degrees of freedom, and so dimensions, to astrocytes computational potential [46, 9, 47]. It is often the case in other modeling studies increasing detail has at worse has no effect, and often leads to improvement [24, 25, 18].

### Predictions

To detect if astrocytes carry on their own computations the first challenge is isolating these signals. [41] has shown this is possible by direct observation. A reasonable default is to say astrocyte activity is a delayed slow filtered reflection of neuronal drive. Under this null any pattern in astrocyte waves is an epiphenomenon of neural activity. If it was then shown that wave patterns are not just reflections, this would be some evidence for astrocyte’s wave computations.

For example, in multi-area, brain-wide recordings the high dimensional dynamics of neurons is well captured by low dimensional dynamical systems [10, 51]. [43] developed convergent cross mapping, a way to compare if two dynamical systems are equivalent using only noisy measurements. This was adapted to work in latent spaces by [7] If astrocyte waves are simple an epiphenomenon this procedure should identify an equivalence between a learned low dimensional model neuronal firing rates, and measurements of astrocytes of calcium dynamics. Taken over several cortical areas, experiments in cross mapping would provide good initial evidence for astrocyte computation and independence.

Another open problem is glia biology is understanding why cortical astrocytes are so spread out. The results for leak and complexity suggests an answer. Astrocytes are spread out to minimize leak, or crosstalk, between themselves. In other words, are they spread out to play a role as waveguides [4, 22]?

## Methods

All learning networks were implemented in python using the pytorch framework. Code and data are available at https://github.com/CoAxLab/glia_playing_atari. Simulations were run on a 4 Nvidia GeForce GTX 1080 Ti cluster.

We trained all models on the XOR problem, and on two computer vision tasks MINST digits and MINST fashion. MINST datasets were sourced from torchvision. Images were randomly assigned to training and reporting sets, and were used as provided. XOR data was its truth table implemented in tensors. Note the XOR problem has no meaningful way to split between training and testing

Neural networks were multilayered perceptrons, with all to all connections. These networks played two roles. The first used a variational autoencoder [30] to reduce input dimensionality into something astrocyte waves can practically learn from. More on this below. The second kind implemented the classification networks, which were the point of comparison for the astrocyte waves. The autoencoder design is discussed in the next section. The hyper-parameters for the three classifier networks are shown in Table 1.

The input image size for both MINST dataset is N by N pixels, which gives flattened vectors of size K. The output size for both is 10, the number of classes to be learned. As astrocytes in simulation needed *l* steps to make a dimensionality change of size *K*. This led to vanishing gradients [17]. To solve this two kinds of dimensionality reduction were considered. One inspired by standard machine learning practice. The use of a variational autoencoder. The other was inspired by the neural circuits, which are often modelled as sparse random projections [13, 3, 6].

To implement astrocyte communication we defined three prototype “steps” for wave propagation. Our use of the term steps is synonymous with “layers” in typical neural networks. We distinguish them because our astrocyte waves may need several steps to accomplish the same output as the equivalent single layer of neurons. If *m* is the number of cells in the input and m the number of cells in the output, we define three step types where *m > n*, *m* = *n*, and where *n < m*. s 1. *Slide* (d) steps have input size *n* and output size *m* = *n*. 2. *Gather* (g) steps have input size *n* and output size max(*n* −2, 1). 3. *Spread* (s) steps have input size *n* and output size *n* + 2. For illustrated examples, see Figure 5.

**Figure 5:**
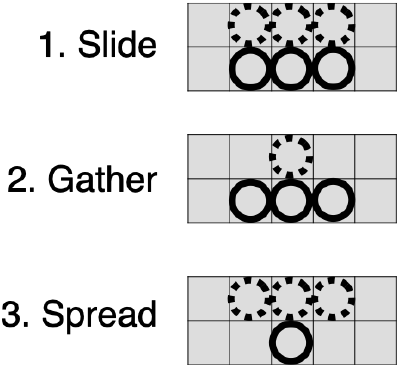
Astrocyte tensor operations – three prototypes for astrocyte steps. Solid circles represent the current cell (the input for that step). Dashed circles represent the projected output (for that step). For compactness in the Tables which describe the networks, I use *s* to denote spread steps, *g* to denote gather, and *d* for slide.

Each step ensures only local, (*i*, *j*, *k*) nearest neighbor interactions. These were implemented as pytorch tensors of full rank *n* and *m* but whose sum operations for each *j*th cell were locally indexed during both forward computation and backpropagation. This meant working around pytorch’s limits on “in place” operations. Each wave step could contain stochastic dropout, or (Gaussian) noise injection or (Gaussian) spatial convolution.

